# MeshMonk: Open-source large-scale intensive 3D phenotyping

**DOI:** 10.1101/491639

**Authors:** Julie D. White, Alejandra Ortega-Castrillón, Harold Matthews, Arslan A. Zaidi, Omid Ekrami, Jonatan Snyders, Yi Fan, Tony Penington, Stefan Van Dongen, Mark D. Shriver, Peter Claes

## Abstract

In the post-genomics era, an emphasis has been placed on disentangling ‘genotype-phenotype’ connections so that the biological basis of complex phenotypes can be understood. However, our ability to efficiently and comprehensively characterize phenotypes lags behind our ability to characterize genomes. Here, we report a toolbox for fast and reproducible high-throughput dense phenotyping of 3D images. Given a target image, a rigid registration is first used to orient a template to the target surface, then the template is transformed further to fit the specific shape of the target using a non-rigid transformation model. As validation, we used N = 41 3D facial images registered with MeshMonk and manually landmarked at 19 locations. We demonstrate that the MeshMonk registration is accurate, with 0.62 mm as the average root mean squared error between the manual and automatic placements and no variation in landmark position or centroid size significantly attributable to landmarking method used. Though validated using 19 landmarks for comparison with traditional methods, MeshMonk allows for automatic dense phenotyping, thus facilitating more comprehensive investigations of 3D shape variation. This expansion opens up an exciting avenue of study in assessing genomic and phenomic data to better understand the genetic contributions to complex morphological traits.

## Introduction

The phenotypic complement to genomics is *phenomics*, which aims to obtain high-throughput and high-dimensional phenotyping in line with our ability to characterize genomes^1^. The paradigm shift is simple and similar to the one made in the Human Genome Project: instead of ‘phenotyping as usual’ or measuring a limited set of simplified features that seem relevant, why not measure it all? In contrast to genomic technologies, which successfully measure and characterize complete genomes, the scientific development of phenomics lags behind. However, with the advent of new technologies, hardware exists for extensively and intensively collecting quantitative phenotypic data. For example, 3D image surface and/or medical scanners provide the optimal means to capture information of biological morphology and appearance. Today, the challenge is to establish standardized and comprehensive phenotypic representations from large scale image data that can be used to study phenotypic variation in the context of genetic variation^2^. This is a challenge that we address with the development of the MeshMonk toolbox, which enables fast and reproducible high-throughput phenotyping of 3D images, or quasi-landmark indication, which can be applied to 3D facial images as well as 3D scans of other complex morphological structures.

Dense correspondence phenotyping is important beyond genomics and could be employed by anthropologists, biologists, and medical clinicians to accurately and reproducibly characterize anatomical structures such that underlying qualities about the structure can be understood. The study of variation and covariation in anatomy can provide insights into the genetic causes and evolution of the anatomical structure. In addition, comparing the anatomy of an individual patient to a control population can indicate pathology to a medical practitioner. Traditionally, this has been achieved using visual clinical assessment or by taking measurements between manually placed anatomical landmarks. Some examples include the endo- and exocanthi (the inner and outer corners of the eyes, respectively) and the pronasale (the tip of the nose).

However, manual landmarking is tedious to perform, difficult to standardize in practice, and prone to intra and inter-operator error^3–7^. Furthermore, sparse landmark configurations can only quantify form at defined landmarks that can be reliably identified and indicated by a human, and thus lack the resolution to fully characterize shape variation in between landmarks. An alternative is to automatically indicate quasi-landmarks across the entire surface of the structure. This is achieved by gradually warping a generic template composed of thousands of points into the shape of each target image through a non-rigid registration algorithm^8–12^. The coordinates of these warped templates, now in the shape of each target, can then be assessed in geometric morphometric analysis. An automatic approach like this is preferable for the analysis of large datasets, avoiding the problems of manual landmarking by multiple operators. Dense phenotype coordinates are also more suitable for applications that require synthesis of a recognizable instance of the actual structure, such as predicting a complete shape from DNA^13^, synthetic growth and ageing of a face^14,15^, constructing 3D facial composites for forensic applications^16^, and characterization of dysmorphology for clinical diagnosis^17,18^.

Surface registration, implemented in MeshMonk, defines a warping of the vertices from one (template) image to their corresponding locations on another (target) image and allows us to quantify and visualize both subtle and acute variation in surface form across a sample by finding the geometrical relationship (one-to-one correspondences) between 3D shapes^8–10,12,19^. The registration strategy is akin to fitting an elastic net onto a solid facial statue through a geometry-driven mapping of anatomically corresponding features. When the template is warped onto each target, the coordinates of any anatomical landmark, manually annotated on the template, can also be defined on each target, thus the complete quasi-landmark indication can also be considered a method for automatic placement of sparse anatomical landmarks^20^. As a validation, we compared a set of 19 sparse landmarks indicated manually by two observers and automatically using MeshMonk.

## Results

### Accuracy

#### Direct comparison of manual and automatic landmark placements

As one measure of validation of the automatic landmark indications, we compared the raw coordinate values of manual landmark indications with the raw coordinate values of automatic landmark indications while considering the manual landmarks to be the “gold standard”. Because of the leave-one-out nature of our approach, we can compare the manual and automatic landmark coordinates directly without fear of training bias. To compare landmark indications, we calculated the root mean squared error between the *x*, *y*, and *z* coordinates for manual and automatic indications (Table 1) and calculated the intraclass correlation coefficient between the *x*, *y*, and *z* coordinates produced by the two methods. When comparing the average of all six manual landmarking indications (C_ML_) and the automatic landmarks trained using this average (C_Auto_), the highest difference after averaging standard deviation values across all axes was 0.85 mm, for the right side exocanthion landmark (Table 1). Overall, the average standard deviation between C_ML_ and C_Auto_ across all landmarks was 0.62 mm. Bland-Altman comparisons showed that the 95% confidence intervals for the landmark indication between methods are within 1.5 mm of a mean difference of 0 mm (Supplementary Fig. S1). Most individuals fall within these confidence limits, with only a few comparisons from each axis having differences greater than 3 mm. The intraclass correlation coefficients for each axis are around 0.99, representing very high correlation and agreement between manual and automatic landmark indications.

**Table 1.**
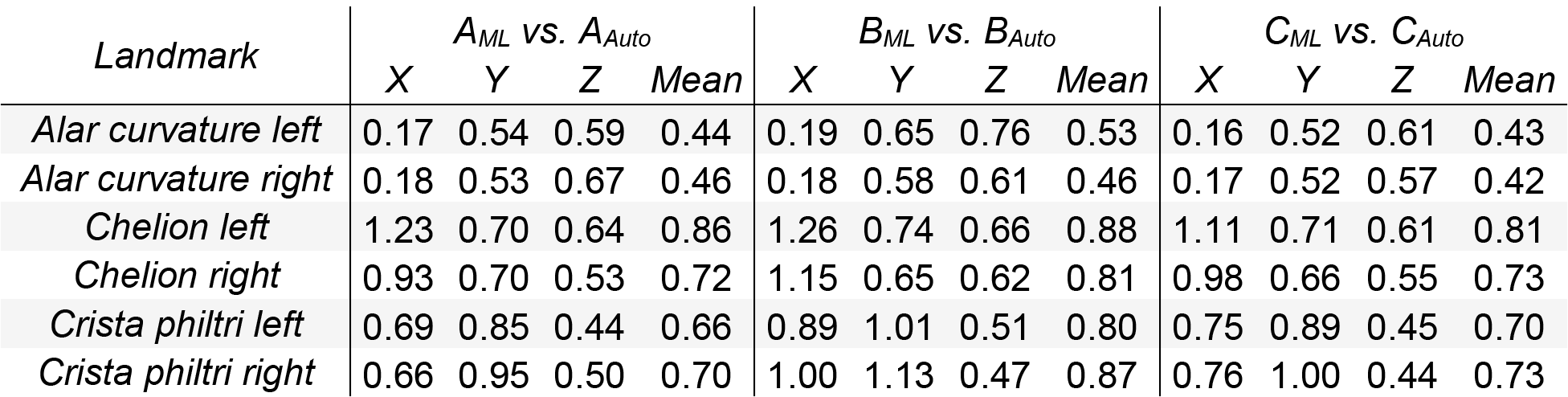
Root mean squared error between manual and automatic landmarks. Root mean squared error (mm) between the manual and automatic landmark indications. Values are presented for each axis, averaged across all faces, as well as averaged across the axes (mean).

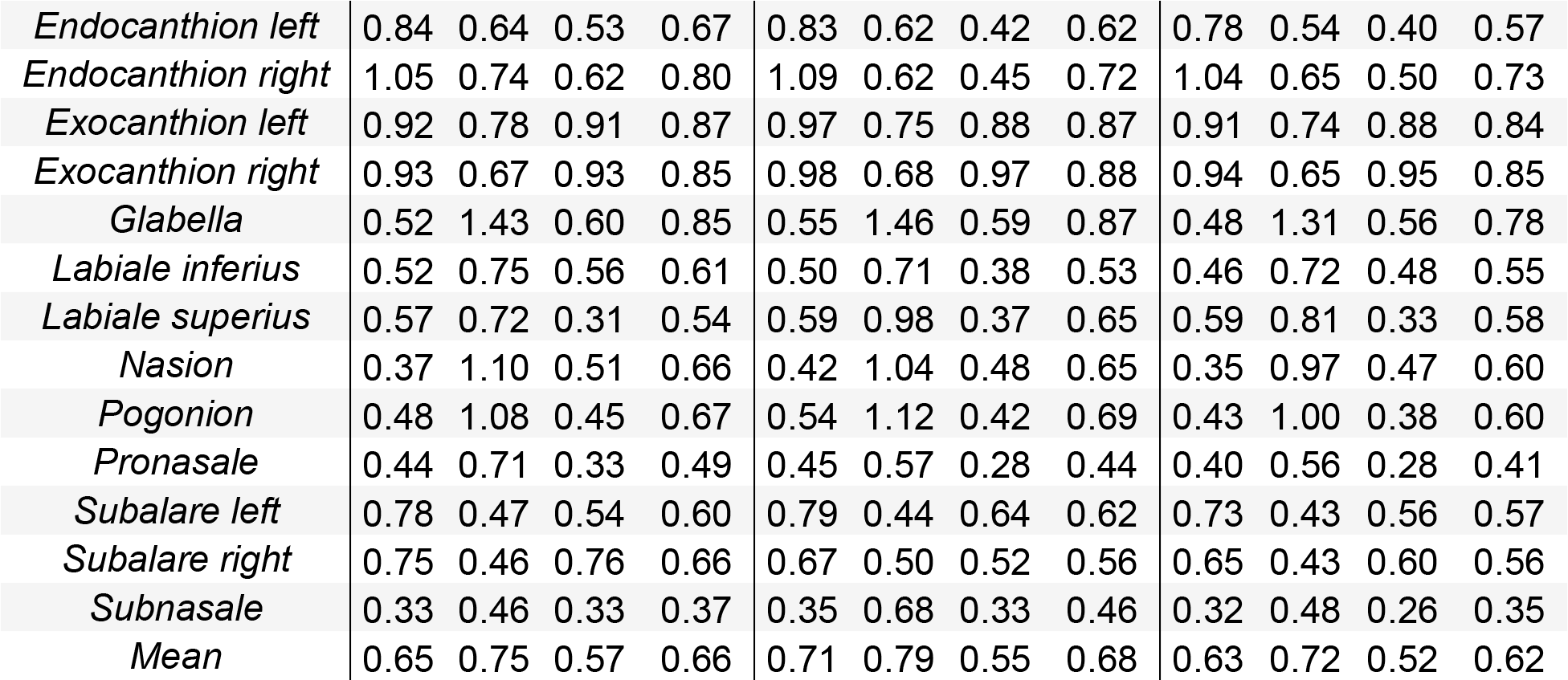

#### Centroid size comparison

We used estimates of centroid size (CS; the square root of the sum of squared distances from each landmark to the geometric center of each landmark configuration) as an additional assessment of the similarity between manual and automatic landmark placements, since centroid sizes feature heavily in geometric morphometric assessments. The ICC of centroid sizes calculated using the manual and automatic landmarks were all high (ICC_A_ = 0.9589, ICC_B_ = 0.9486, ICC_C_ = 0.9591; Supplementary Fig. S2). Analysis of variance (ANOVA) by individual, observer, and method shows that individual is the only significant factor in explaining variance in centroid size (*F* = 130.407, *p* < 2 × 10^−16^; Table 2). Bland-Altman comparison showed that the 95% confidence intervals for the centroid size estimates between methods are 2 mm relative to an average centroid size of about 165 mm (Supplementary Fig. S2).

**Table 2.**
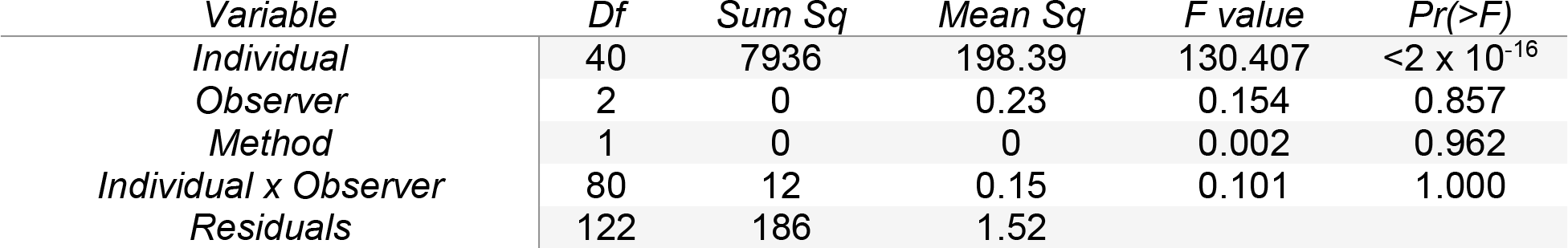
ANOVA of centroid sizes. Results from an ANOVA with centroid size as the response variable and individual, observer, method and individual x observer as predictors.

| *Variable* | *Df* | *Sum Sq* | *Mean Sq* | *F value* | *Pr(>F)* |
| --- | --- | --- | --- | --- | --- |
| *Individual* | 40 | 7936 | 198.39 | 130.407 | $<2 \times 10^{-16}$ |
| *Observer* | 2 | 0 | 0.23 | 0.154 | 0.857 |
| *Method* | 1 | 0 | 0 | 0.002 | 0.962 |
| *Individual x Observer* | 80 | 12 | 0.15 | 0.101 | 1.000 |
| *Residuals* | 122 | 186 | 1.52 |  |  |

#### Analysis of shape variance

A multivariate analysis of variance (MANOVA) on shape, based on the average of each observer’s manual landmark indications and automatic landmark configurations, separately, was performed to determine if the variance explained by individual and observer factors was similar in both methods (Supplementary Table S1). In both methods, individual variation contributed to most of the variation in shape (R^2^_ML_ = 94%; R^2^_Auto_ = 97%). Differences in observer accounted for 1.9% of the variation in shape from manual landmarks and 2.6% of the variation in shape from automatic landmarks. In total, 3.9% of the variation present in manual landmark shape configurations was unexplained by our model while only 0.22% of the variation was unexplained when testing the automatic landmark configurations. A MANOVA on Generalized Procrustes Analysis (GPA) aligned manual and automatic configurations from each observer, with method, individual, observer, and individual x observer as predictors showed that landmarking method did not significantly account for variation in landmark placement (*F* = 0.3463; *p* = 0.987; Table 3).

**Table 3.** MANOVA on manual and automatic landmarks. Results from a single MANOVA 624 using the average manual landmark indications from each observer (A_ML_ and B_ML_) and the 625 automatic landmark indications using the observer level averages (A_Auto_ and B_Auto_).

| *Variable* | *DF* | *SS* | *MS* | *R<sup>2</sup>* | *F* | *Z* | *Pr(>F)* |
| --- | --- | --- | --- | --- | --- | --- | --- |
| *Method* | 1 | 0.0003 | 0.0003 | 0.0004 | 0.3463 | -2.2135 | 0.987 |
| *Individual* | 40 | 0.6522 | 0.0163 | 0.8778 | 20.2019 | 23.3507 | 0.001 |
| *Observer* | 1 | 0.0167 | 0.0167 | 0.0224 | 20.6396 | 11.4067 | 0.001 |
| *Individual x Observer* | 40 | 0.0085 | 0.0002 | 0.0114 | 0.2623 | 13.7253 | 0.001 |
| *Residuals* | 81 | 0.0654 | 0.0008 | 0.0880 |  |  |  |
| *Total* | 163 | 0.7430 |  |  |  |  |  |

### Reliability

#### Intra- and inter-observer error of manual landmarks

The quantitative study of morphology using 3D coordinates requires specific attention to measurement error and has a robust presence in the literature. For each observer, we calculated the intra-observer error of the manual landmarks as the standard deviation between the *x*, *y*, and *z* coordinates of each observer’s three landmarking iterations. Supplementary Table S2 reports intra-observer standard deviations for the manual landmark indications along each axis, averaged across images. The average standard deviation of observer A across all landmarks was 0.58 mm while the average standard deviation of observer B across all landmarks was 0.44 mm. The average inter-observer error, measured as the standard deviation between the average *x*, *y*, and *z* coordinates of each observer’s landmarking iterations was 0.40 mm. This range of deviation is considered highly precise and is similar to previously reported measures of landmark error^6,21^.

The analysis of measurement and observer error for the manual landmarks alone, assessed using a MANOVA for shape, with individual, observer, observer x individual, and nested observer x landmarking iteration as factors showed that non-individual factors contributed significantly to variation in shape (Supplementary Table S3). Individual variation contributed to most of the variation in shape (85%), as expected. Simple measurement error accounted for 3.5% of the total variation in shape. Additional to this, differences in observer accounted for 1.8% of shape variation, and deviation across landmarking iterations contributed an additional 1.5% of the total variation in shape. In total, non-individual effects contributed to 15% of the total shape variation, with 8.3% of this variation unexplained by the model.

#### Comparison of manual and automatic inter-observer errors

By treating the automatic landmark indications as if they were performed by a third observer, we calculated “inter-observer” errors to compare the variation of automatic and manual landmarking. In this assessment, we compared inter-observer errors calculated using only the manual landmarks (A_ML_ vs. B_ML_) with error estimates calculated by replacing one of the observer’s manual landmark indications with the automatic indications trained using that observer’s average. This resulted in two extra estimations of inter-observer error (A_ML_ vs. B_Auto_ and A_Auto_ vs. B_ML_), calculated as the standard deviation between *x*, *y*, and *z* coordinates (Supplementary Table S4; Supplementary Fig. S3). The mean manual landmarking inter-observer error was 0.40 mm while both manual-automatic comparisons had mean standard deviation values of 0.53 mm (Supplementary Table S4). A paired t-test between the manual landmark error values and each of the manual-automatic comparison showed that the landmark indications that were significantly different between the two methods tended to be those where facial texture likely assisted in the placement of the manual landmarks (e.g. localizing the crista philtra by looking at the differences in color between the lips and the skin; Supplementary Table S5). This result indicates that automatic sparse landmarking using MeshMonk will likely produce more robust results when given input data that has a strong anatomical orientation (e.g. the nasion and pogonion). Even given these differences in variance, the manual-automatic comparisons did not produce errors that were completely outside the range of inter-observer errors, a sign of the reliability of the MeshMonk registration.

As an illustration of the low errors between automatic landmark indications trained using different observers, we calculated the standard deviation between automatic landmark indications trained using the average of observer A’s three landmark indications and the average of observer B’s three landmark indications (A_Auto_ vs. B_Auto_; Supplementary Table S6, Supplementary Fig. S3). The variance of the average standard deviation values were significantly different for all landmarks except labiale superius, where we could not reject the null hypothesis that the variances of the two standard deviation distributions were equal (*F* = 2.4213, *p* = 0.1236). Supplementary Fig. S3 shows that the variance between automatic landmarking indications (A_Auto_ vs. B_Auto_) is easily identified as being smaller than the manual landmark inter-observer error (A_ML_ vs. B_ML_).

## Discussion

Through studies utilizing manually placed sparse landmarks, we have begun to understand the biological basis and evolution of complex phenotypes, both normative and clinical. However, there is still much to be learned. One avenue for improvement is to expand and speed up the production and analysis of data using methods derived from engineering and computer vision, which allow for the description of shapes as “big data” structures instead of sparse sets of landmarks or linear distances, thus matching our ability to describe phenotypes with our ability to describe genomes. To this end, we introduce the MeshMonk registration framework, giving researchers the opportunity to quickly and reliably establish a homologous set of positions across entire samples. We have validated this framework using a sparse set of landmarks, though the registration framework produces thousands of landmarks to finely characterize the structure.

MeshMonk represents a step forward in our ability to describe complex structures, like the human face, for clinical and non-clinical purposes. Consider Figure 1, showing the starting template for facial image registration (left) as well as three example faces (right). Each point on the images represents a quasi-landmark data point that is homologous and can be compared across faces. Researchers are no longer limited to a few homologous points, chosen because they can be reliably indicated over hundreds of hours of work. Instead, minute details of the face can be identified and compared across thousands of images in a few hours, and additional images can be incorporated just as easily, regardless of the camera system with which they were captured, allowing for the incorporation of images from different sources and databases (e.g. Facebase.org).

**Figure 1.**
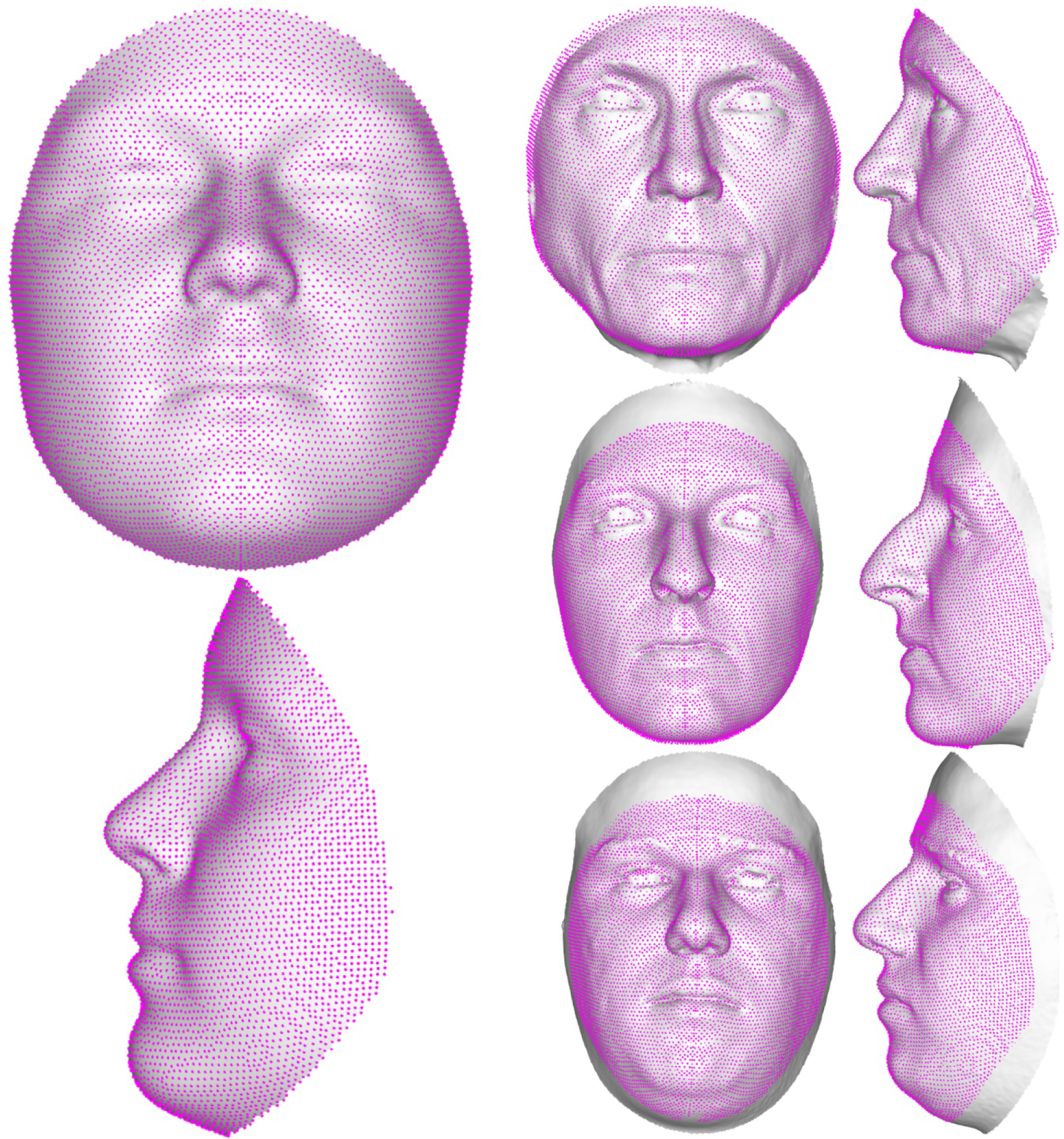
Facial template registration. The template (left), built as the average of more than 8000 admixed facial scans, can easily wrap onto any face (three example faces on the right), accurately representing its particular traits. This allows for the explanation of any face in the template’s coordinates, enabling a spatially-dense analysis between any registered surfaces.

Because of the relative newness of dense correspondence phenotyping, few studies have focused on the accuracy and reliability of the resulting registrations. Previous studies using versions of the MeshMonk framework have shown that the error associated with the registration of the template onto facial images is 0.2 mm^22^ and parameters of the toolbox have been fine-tuned, as discussed elsewhere^8^ and in the Supplemental Methods. To provide some validation regarding the ability of the registration process to accurately identify anatomical positions of interest, we used a set of 40 faces with manual landmark indications to “train” positions of interest on the template, then automatically indicate these positions on a face that was not present in the training dataset. In the comparison of manual and automatic landmark indications, the positions of the manual landmarks were considered to be the gold standard, as they have a long history of use and validation in morphological studies^5,21^. By limiting ourselves to a set number of sparse landmarks, we cannot necessarily speak to the accuracy of structures not involved in our validation (i.e. the cheeks), but we argue that the results for our comparison speak highly of the fidelity with which the MeshMonk registration framework aligns to underlying anatomical structures.

In the direct comparison of sparse landmarks placed manually and using the MeshMonk toolbox, the average difference between the manual and automatic placements was low (Supplementary Fig. S1), with the average root mean squared error across all landmarks ranging from 0.62 to 0.68 mm (Table 1), which is well within the range of acceptable error for manual landmarks^5,6,21^ and similar or below errors reported in other comparisons of manual and automatic landmarking methods^23–26^. When assessing landmarking methods separately, the variance in landmark configuration attributable to individual and observer factors is similar, with considerably less variation left unexplained by a MANOVA model using automatic landmark configuration as the response (Supplementary Table S1). When assessing manual and automatic landmark configurations in a single MANOVA, the landmarking method is a nonsignificant factor, indicating that variation in scans is not attributable to variation in landmarking method (Table 3). This result was also reproduced when comparing centroid sizes calculated using manual and automatically placed landmarks (Table 2), speaking to the high correspondence between landmark indications placed by human observers and those indicated by the MeshMonk toolbox.

The validation results together suggest that the MeshMonk toolbox is able to reliably reproduce information given by manual landmarking. Though the larger contribution of the MeshMonk toolbox is the ability to quickly and densely characterize entire 3D surfaces, our illustration using a small number of manually placed landmarks as a training set could be useful for studies seeking specifically to study a sparse set of landmarks, perhaps to add more images to a dataset that is already manually landmarked or to add additional landmarks to an analysis. Utilization of the MeshMonk toolbox also gives the opportunity to minimize variation due to different observers. Take, for example, datasets with manual landmarks indicated by two different observers. During the course of analysis, the inter-observer error of these observers would have to be calculated and taken into account when interpreting results. From our own study, the inter-observer error of the manual landmarks placed by two different observers was 0.40 mm (Supplementary Table S2). With the automatic landmarking framework implemented during this study, we can minimize both intra-observer variance for a single scan (by averaging together all indications of that scan by a single observer) and intra-observer variance across scans by placing all indications from the training dataset on the template mesh and averaging the entire training set before using MeshMonk to place them in an automatic fashion on the target image. This process finely tunes the position of the landmark, such that even if the training sets were indicated by two different observers, the variation in automatic landmark indication is much smaller than the variation in manual landmark indication, averaging 0.27 mm in our study (Supplementary Table S6; Supplementary Fig. S3).

A visual hallmark of the ability of spatially dense surface registration to reliably represent anatomical structures is found in the crispness of “average shapes,” constructed by averaging together all registered surfaces in a study sample. Because the MeshMonk registration aligns closely with the underlying anatomical structure, averages across the study samples continue to cleanly resemble the structure and detail is not lost in the averaging process. As depicted in Figure 2, consider the sample average of the 41 faces in this work and 100 mandible scans. In the rigid-only averages, details are overly smoothed compared to the level of detail present in the rigid plus non-rigid registration averages. For example, it is obvious to the naked eye that the sharpness of the eyes, nose, philtrum and mouth for the facial average, and the alveolar crest, mental foramen, and coronoid and condylar processes for the mandible, are clearly better represented with the rigid plus non-rigid registration. Thus, non-expert readers can easily evaluate the quality of dense-correspondence morphometrics research by looking at the average surfaces, which are typically used in manuscript figures, with the understanding that high quality registration leads to sharp average scans where anatomical positions of interest are clearly defined.

**Figure 2.**
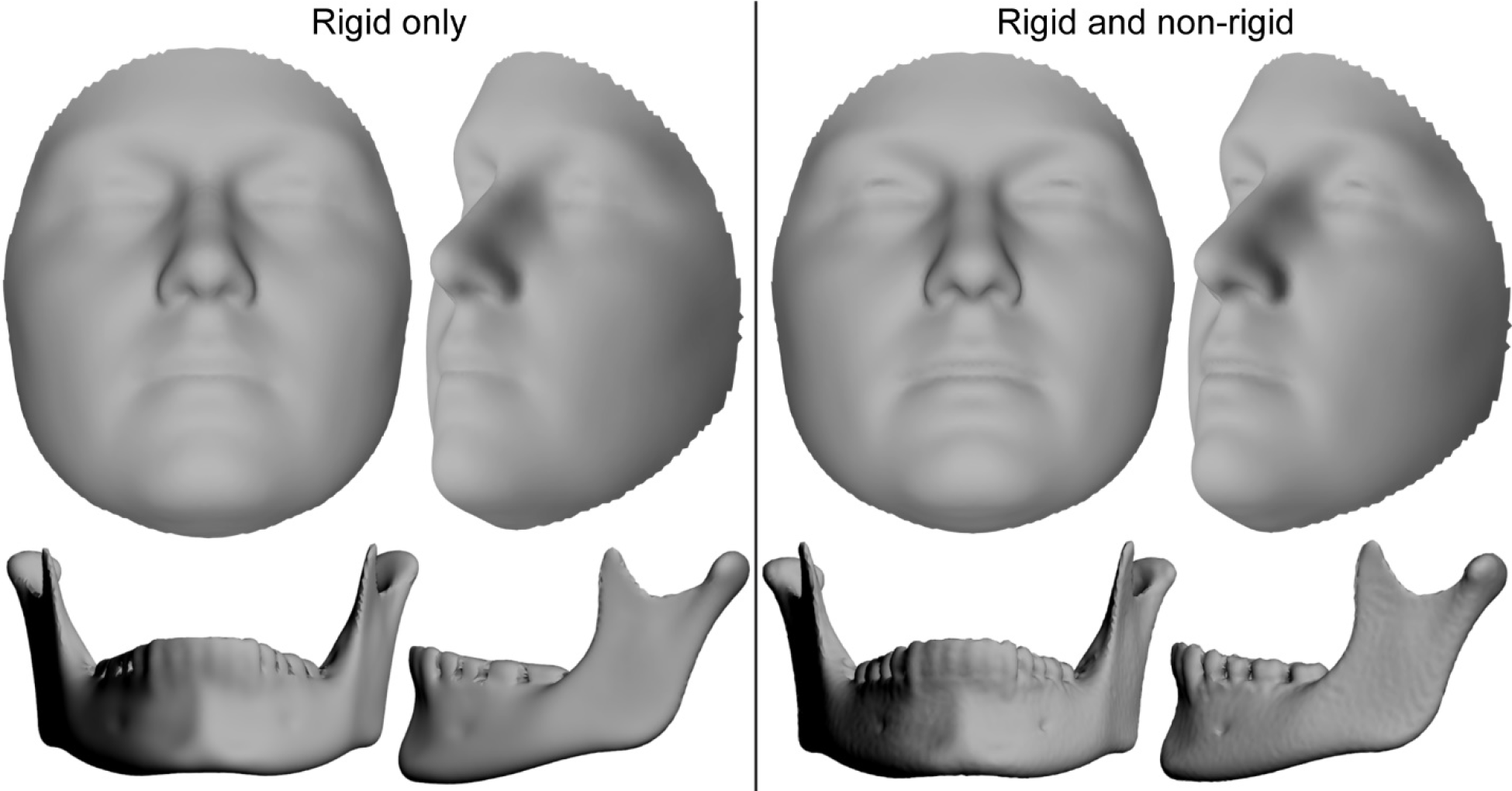
Comparison of rigid and non-rigid registration algorithms. Sample averages using the 41 validation faces and 100 mandible scans. Scans were registered using rigid registration only (left) and then simply mapped exactly to their closest point on the target surfaces or mapped using rigid plus non-rigid (visco-elastic) registration (right).

In this study, we present MeshMonk, an open-source resource for intensive 3D phenotyping on a large scale. Through dense-correspondence registration algorithms, like MeshMonk, we can advance our ability to integrate genomic and phenomic data to explore variation in complex morphological traits and answer evolutionary and clinical questions about normal-range variation, growth and development, dysmorphology, and taxonomic classification.

## Materials and Methods

### Explanation of Meshmonk registration

The core functionality of the MeshMonk toolbox is implemented in C++, with a focus on computational speed and memory to enable the processing of large datasets of 3D images. Interaction with the toolbox is provided using MATLAB™, enabling an easy to use implementation and visualization environment for the user. A schematic of the complete surface registration algorithm is presented in Figure 3 and a short video of the registration on this example face is also available at the following GitHub account (https://github.com/juliedwhite/MeshMonkValidation/).

**Figure 3.**
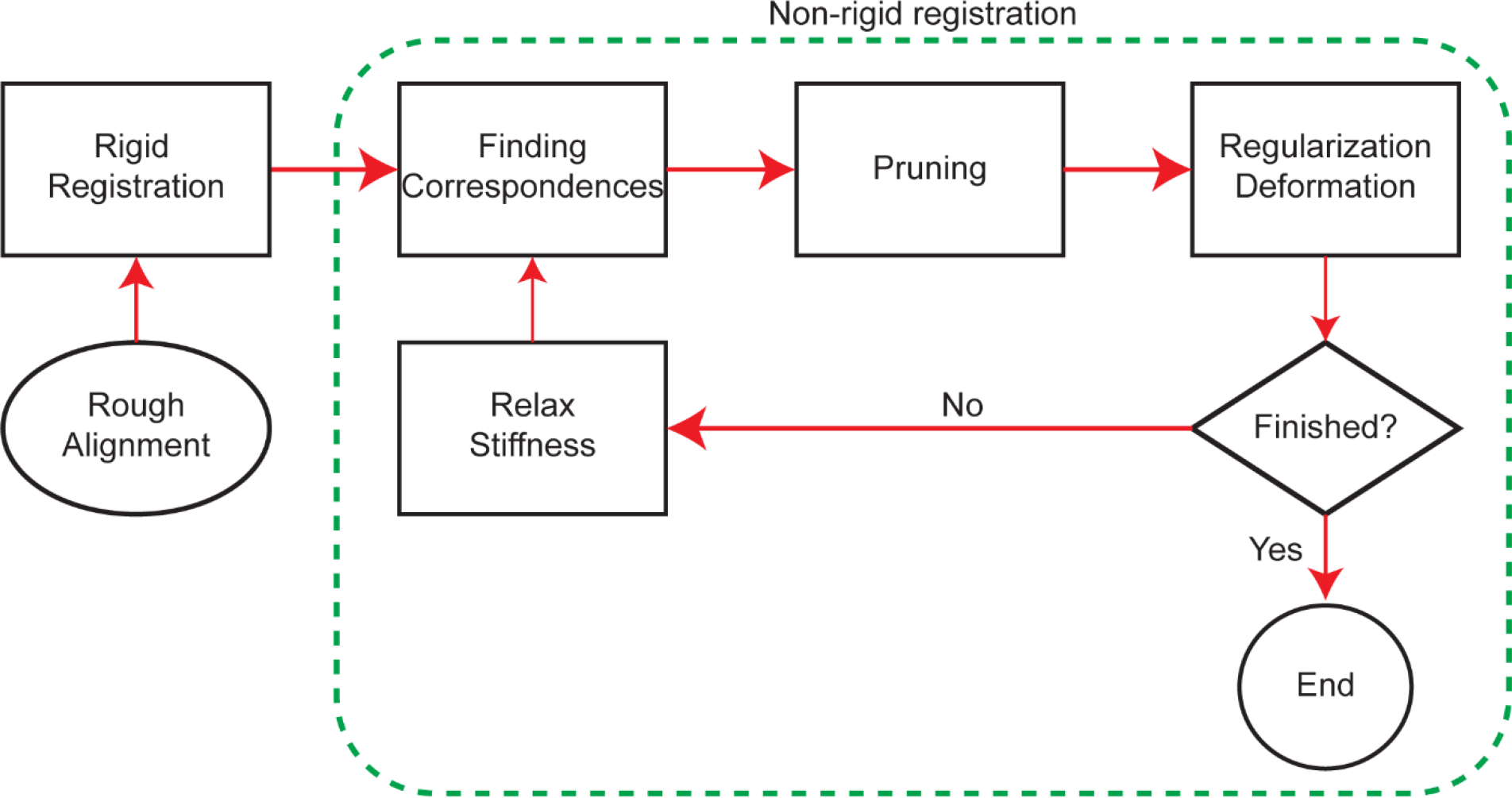
Schematic of the MeshMonk’s surface registration algorithm. MeshMonk uses an initial rigid registration based on the ICP algorithm. This step might require an initial rough alignment to ensure similar orientation, which can be done by placing few landmarks on the target surface. Then, the symmetrical weighted k-neighbor correspondences are found, and outliers are detected and removed. Finally, the visco-elastic transformation is applied. This is performed in an iterative manner, until either a pre-set number of iterations or a pre-set amount of coverage (e.g. a pre-defined root mean squared distance of all template points to the target surface after the transformation) has been reached. Otherwise, the correspondences are updated and the non-rigid registration starts over.

To initiate the process, a rigid registration based on the iterative closest point algorithm^27^ is performed to better align the template to the target surface. During the rigid registration, the transformation model is constrained to changing the position (translation), orientation (rotation), and scale of the template only. Subsequently, a non-rigid registration is done that will alter the shape of the template to match the shape of the target surface. During the non-rigid registration, a visco-elastic model is enforced, ensuring that points that lie close to each other move coherently^8^. At any iteration during the registration, for both the rigid and non-rigid registration steps, correspondences are updated by using pull-and-push forces (symmetrical correspondences)^28^ and a weighted k-neighbor approach (Supplementary Fig. S4)^8^.

3D surface images typically contain artifacts such as holes and large triangles indicating badly captured or missing parts. Any correspondence to such artifacts is meaningless and are indicated as correspondence outliers, not to be considered when updating the transformation model, though they are consistently transformed along with the inliers. The MeshMonk toolbox allows for the identification of outliers either deterministically or stochastically, or a combination of both. In each iteration, correspondences are updated and outliers are identified, then an updated transformation model is used. The smoothness of the transformation model is parametrized by convolving the displacement vectors between corresponding points with a Gaussian^29^. The amount of smoothing is high (multiple Gaussian convolution runs) at the beginning iterations, when correspondences are still noisy and hard to define, and reduces gradually towards the later iterations, when correspondences are more accurately defined.

### Parameters and tuning

Given a dataset of 3D images of interest, the entire MeshMonk procedure can be optimized by setting a variety of parameters in the toolbox, and a parameter tuning can be done based on two “quality” measures. First, a quality of “shape fit” is defined as the root mean squared distance of all template points to the target surface after registration. This essentially measures how well the shape of the template was adapted to the target shape and can be measured over multiple images to deduct an overall quality of shape fit from the dataset. Second, an indication of the consistency of point indications across the same dataset is obtained following the principle of minimum description length in shape modelling^30^. Given two models explaining the same amount of variance, the model requiring fewer parameters is favored, or given two models with the same number of parameters, the one explaining more variance in the data is favored. To this end, a principal component analysis (PCA) is used to assess registration quality because, if the point indications were performed consistently, few PCs are required to explain variation in the registration results. A parameter tuning was done for the facial data in this work prior to the validation and is described in the Supplemental Methods.

### Validation sample and data curation

Our collaborative group has recruited participants through several studies at Pennsylvania State University, recruited at the following locations: State College, PA (IRB 44929 and 4320); New York, NY (IRB 45727); Urbana-Champaign, IL (IRB 13103); Dublin, Ireland; Rome, Italy; Warsaw, Poland; and Porto, Portugal (IRB 32341). All procedures were performed in accordance with the relevant guidelines and regulations from the Pennsylvania State University Institutional Review Board and all participants signed a written informed consent form before participation. Participants additionally gave optional informed consent for publication of their images in a variety of formats, including online open-access publications.

Stereo photogrammetry was used to capture 3D facial surfaces of N~6,000 participants using the 3dMD Face 2-pod and 3-pod systems (3dMD, Atlanta, GA). This well-established method generates a dense 3D point cloud representing the surface geometry of the face from multiple 2D images with overlapping fields of view. During photo capture, participants were asked to adopt a neutral facial expression with their mouth closed and to gaze forward, following standard facial image acquisition protocols^31^.

### Manual placement of validation landmarks

Of the larger sample, N=41 surface images were chosen at random for validation, excluding participants who reported facial surgery or injury. These images were diverse with respect to sex, age, height, weight, and 3D camera system used (Supplementary Table S7). 3dMDpatient was used to record the 3D coordinates of 19 standard landmarks (7 midline and 12 bilateral) from each unaltered surface in wavefront.obj format (Supplementary Fig. S5; Supplementary Table S8). Two independent observers placed landmarks three times each, with at least 24 hours in-between landmarking sessions, resulting in six total landmark indications for each facial image. For each individual, we checked for gross landmark coordinate errors before analysis. In the subsequent analysis, A_ML_ represents the average manual landmarks from observer A, B_ML_ represents the average manual landmarks from observer B, while the combined average of all six manual landmark indications is denoted as C_ML_.

### Automatic placement of validation landmarks

To obtain automatic indications of the 19 validation landmarks, each of the validation faces was registered using MeshMonk and the manual landmark placements were transferred to the registered face by coordinate conversion (Figure 4A)^32^. Because the registered faces are now in the same coordinate system as the original template, we can subsequently transfer the manual landmark indications to the original pre-registration template, giving a set of 41 × 2 observers × 3 indications = 246 manual landmark positions on the template scan (Figure 4B). One by one, each face was left out while averaging the other 40 landmark placements to “train” the automatic landmarks (Figure 4C). These averages were then transferred onto the left-out (target) face, resulting in the automatic placement of the validation landmarks using a “training” set that did not include the target face (Figure 4D). Further detail on this process can be found in the Supplemental Methods.

**Figure 4.**
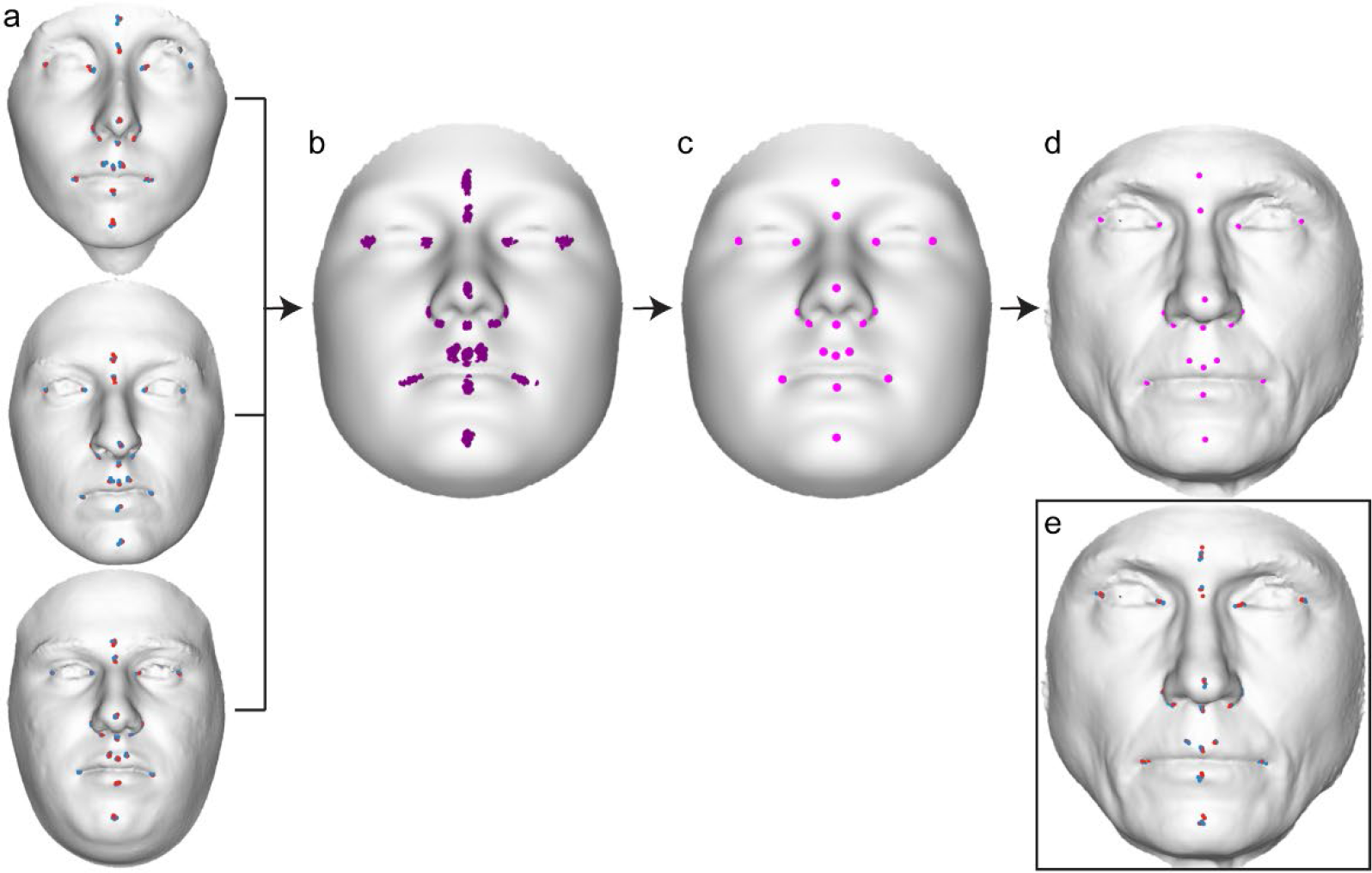
Depiction of automatic landmark indication. **(a)** Each facial scan was manually landmarked six times, three times each by two observers (red and blue points). **(b)** These iterations were then averaged together and are placed on the template (purple points). **(c)** The average of all but the test face (N=40) placements on the template, serving as the foundation for the automatic landmark placements (magenta points). **(d)** Coordinate conversions, described in more detail in the Supplemental Methods, is used to subsequently transfer the automatic landmark placements from the template to the target (left-out) surface, serving as the automatic landmark indication for the target surface (magenta points). **(e)** The manual landmark indications from two observers (red and blue points) for the shown example face, for comparison to the automatic indication in (d).

The placement of automatic landmarks was performed three times: once using the average of observer A’s manual landmark indications as input (A_Auto_), again using the average of observer B’s manual landmark indications (B_Auto_), and a final time using the combined average of all six manual landmark indications from both observers (C_Auto_). This process resulted in three placements of automatic landmarks for comparison.

### Accuracy

We assessed the accuracy of the MeshMonk automatic landmark placements by calculating the root mean squared error (RMSE) between manual and automatic coordinates. We also calculated Bland-Altman^33^ and Intraclass Correlation Coefficient (ICC)^34^ statistics to compare the manual and automatic landmark indications. The Bland-Altman method is preferred over correlation or regression as it is less influenced by the variance of the sample and ICC is preferred because it tests both the degree of correlation and agreement between methods. We additionally compared estimates of centroid size calculated using each method and performed an ANOVA on the centroid size calculations, with individual, observer, method, and individual x observer as predictors, to determine if variation in centroid size could be attributable to variation in landmarking method.

We utilized several methods to determine if the variance structures produced by the two methods were similar. Fitting a MANOVA estimates the variance explained, in correlated outcome variables, by various factors included in the model. Here, we performed MANOVAs separately on the GPA-aligned average manual landmark indications from each observer (A_ML_ and B_ML_) as well as on the GPA-aligned automatic landmark indications trained using the average of each observer’s three landmark placements (A_Auto_ and B_Auto_), with image and observer as predictors in both tests. By comparing the results of these two tests, we can determine how the explanation of shape variance changes given a different landmarking method. To directly determine if any variance in shape was attributable to landmarking method, we combined the average manual landmark placements of each observer with the automatic placements trained using each of these averages and aligned them using GPA (A_ML_, B_ML_, A_Auto_, and B_Auto_). We then tested the shape variation in this combined space as the response in a MANOVA, with individual, observer, method, and individual x observer as factors.

### Reliability

We calculated the manual landmarking intra-observer error, the variation between indications taken at different times by the same individual, as the standard deviation between the *x*, *y*, and *z* coordinates of each observer’s manual landmarking indications. The inter-observer error, the difference between manual landmark indications made by different individuals, was calculated as the standard deviation between each observer’s average *x*, *y*, and *z* coordinates (A_ML_ vs. B_ML_). As an additional method to understand the variation present in the manual landmark indications only, we performed a MANOVA after GPA-aligning the six manual landmarking indications^35^. Study individual, observer, and landmarking iteration were used as factors and landmark configuration as the response.

To determine if the automatic indication process was more or less variable than manual landmarking, we compared the inter-observer error calculated using only the manual landmarks (A_ML_ vs. B_ML_) to the standard deviation between one observer’s manual landmarks and the automatic landmarks trained using the other observer’s manual placements (A_ML_ vs. B_Auto_ and A_Auto_ vs. B_ML_), as if the automatic indications replaced the manual indications in a calculation of inter-observer error. A paired T-test was used to determine whether the “inter-observer errors” calculated using the automatic indications were significantly different than the error calculated using only the manual indications. Standard deviation values calculated using both automatic placements (A_Auto_ vs. B_Auto_) were compared to manual landmarking inter-observer error to illustrate the variance of automatic landmark indications. Levene’s test^36^ was performed to determine if the variances of the inter-observer errors calculated using the manual landmarks were equal to the standard deviation between the automatic landmarks (the null hypothesis). Levene’s test was chosen because the distribution of standard deviation values was non-normal.

All validation analyses were performed in R using the Geomorph^37^, BlandAltmanLeh (https://cran.r-project.org/web/packages/BlandAltmanLeh/BlandAltmanLeh.pdf), and ICC (https://cran.r-project.org/web/packages/ICC/ICC.pdf) packages, as well as packages for data manipulation (readxl, reshape2, plyr, car, data.table, dplyr, broom) and graphing (ggplot2, GGally, GGpubr). Centroid sizes were calculated using Geomorph and MANOVAs for shape variation were implemented using the ProcD.lm function from Geomorph^37,38^.

## Data Availability Statement

The informed consent with which the data were collected does not allow for dissemination of identifiable data to persons not listed as researchers on the IRB protocol. Thus, the full surface 3D facial images used for validation cannot be made publicly available. In the interest of reproducibility, we have provided the 19 manual and automatic landmarks used for validation as well as the code used to analyze them. These data are available in the following GitHub repository: https://github.com/juliedwhite/MeshMonkValidation/. The MeshMonk code and tutorials are available at https://github.com/TheWebMonks/meshmonk.

## Acknowledgments

MeshMonk was developed with WebMonks (https://webmonks.vision), a Belgian startup that works as a bridge to highly qualified developers in third-world countries, and we are very grateful for their high-quality implementation and support. We also thank the many participants who have volunteered their time and all the past and present members of the Shriver and Claes labs, without whom we would have never been able to develop this toolbox or perform the research that it has contributed to.

## Author Contributions

JW performed all landmark based analyses and landmarked the 3D scans used for validation with AZ. PC and AO performed the parameter tuning on the facial data and provided the automatic landmark indications. JW, AO, and HM wrote the first draft of the manuscript under supervision of PC. HM, YF, and TP provided input and images using mandible scans. PC and JW conceptualized the design of the study. OE, SV, and MS provided input throughout the analyses and writing process. JS developed the MeshMonk code.

## Conflict of Interest

Author J. Snyders is employed by the Belgian non-profit company WebMonks. All other authors declare that the research was conducted in the absence of any commercial or financial relationships that could be construed as a potential conflict of interest.

## Funding

The sample collection and personnel involved in this work was supported by grants from the US National Institute of Justice (2008-DN-BX-K125), the US Department of Defense, the University of Illinois Interdisciplinary Innovation Initiative Research Grant, the Science Foundation of Ireland Walton Fellowship (04.W4/B643), the Penn State Center for Human Evolution and Development, the Research Fund KU Leuven (BOF-C1, C14/15/081) and the Research Program of the Fund for Scientific Research - Flanders (Belgium) (FWO, G078518N).

