## Supplemental Information for "MeshMonk: Open-source large-scale intensive 3D phenotyping"

### *Supplementary Figures*

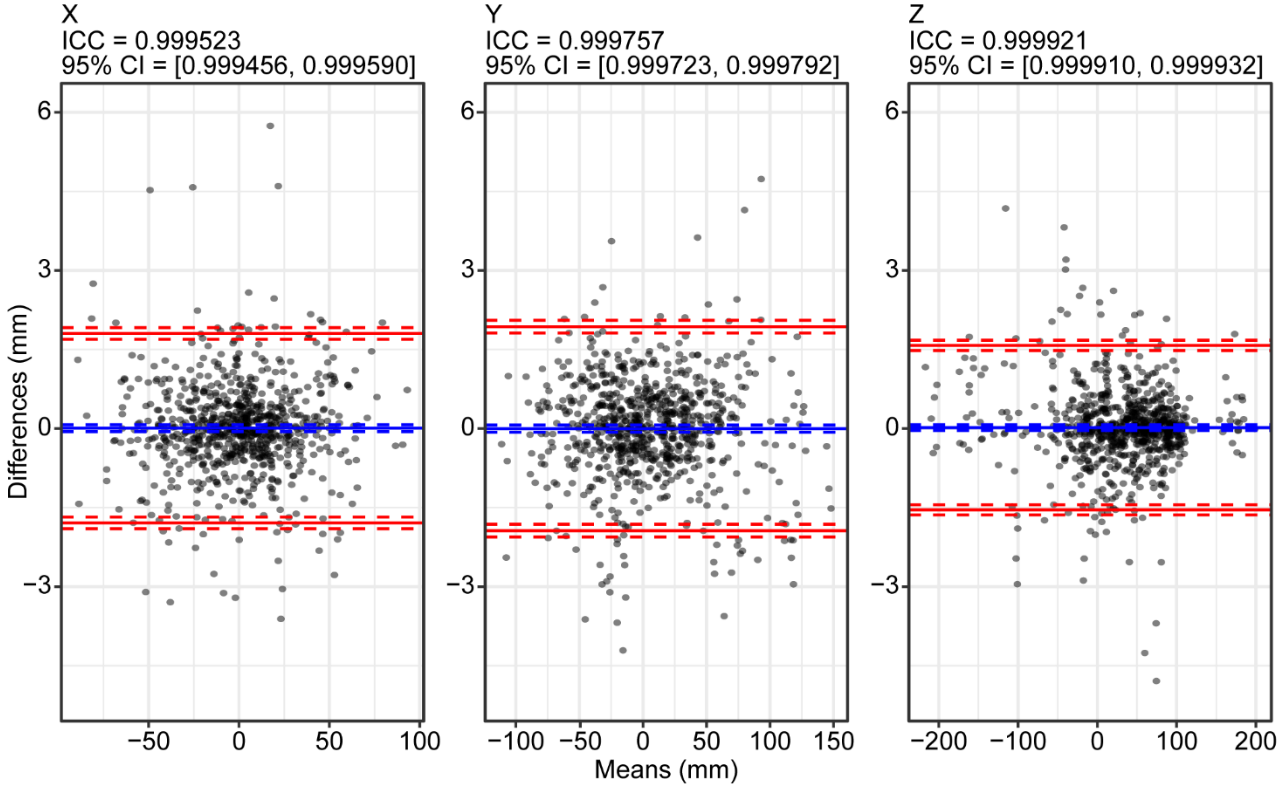


**Supplementary Figure S1. Bland-Altman plot for similarity between manual and automatic landmark placements**. For x, y, and z coordinates, Bland-Altman plot showing the differences between the manual ($C_{ML}$) and automatic ($C_{Auto}$) landmark indications against the averages of the two techniques. Blue lines represent the mean difference value and red lines represent the upper and lower 95% confidence limits. Also given are the intra-class correlation coefficient with ICC 95% confidence interval.

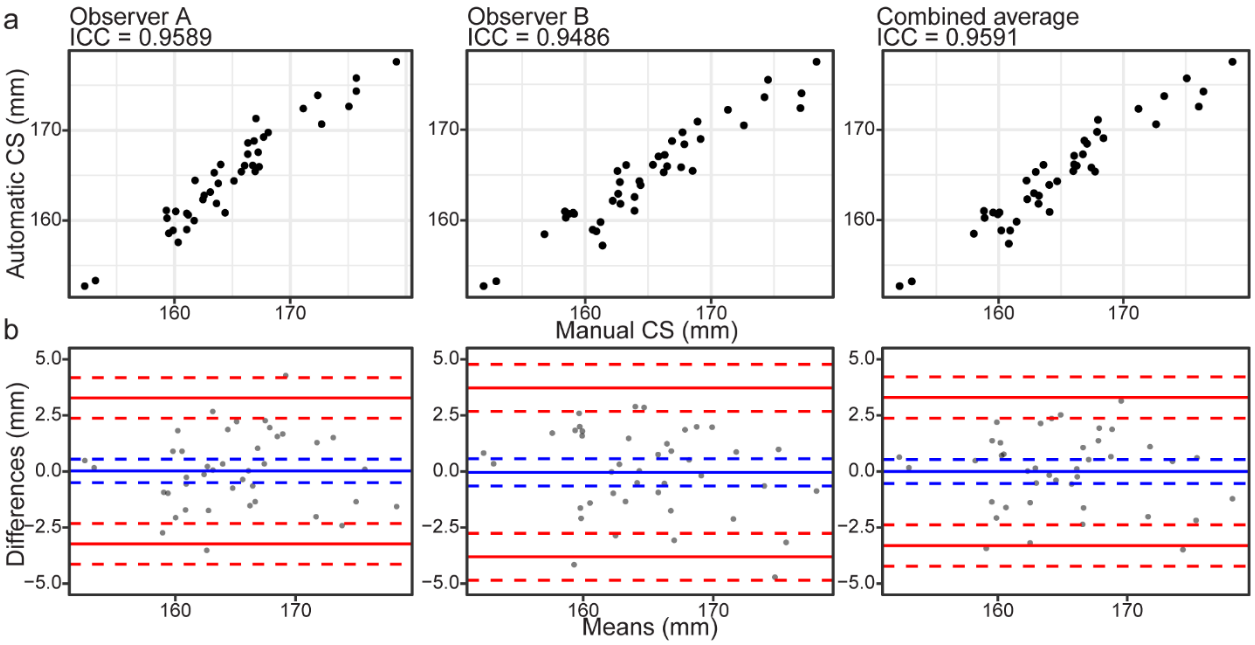


**Supplementary Figure S2. Comparison of centroid sizes.** (**a**) Point plots for comparison of centroid sizes using automatic and manual landmarking methods, separated by observer. (**b**) Bland-Altman plot showing the differences between centroid sizes produced using the manual and automatic methods against the averages of the two techniques. Blue lines represent the mean difference value and red lines represent the upper and lower 95% confidence limits.

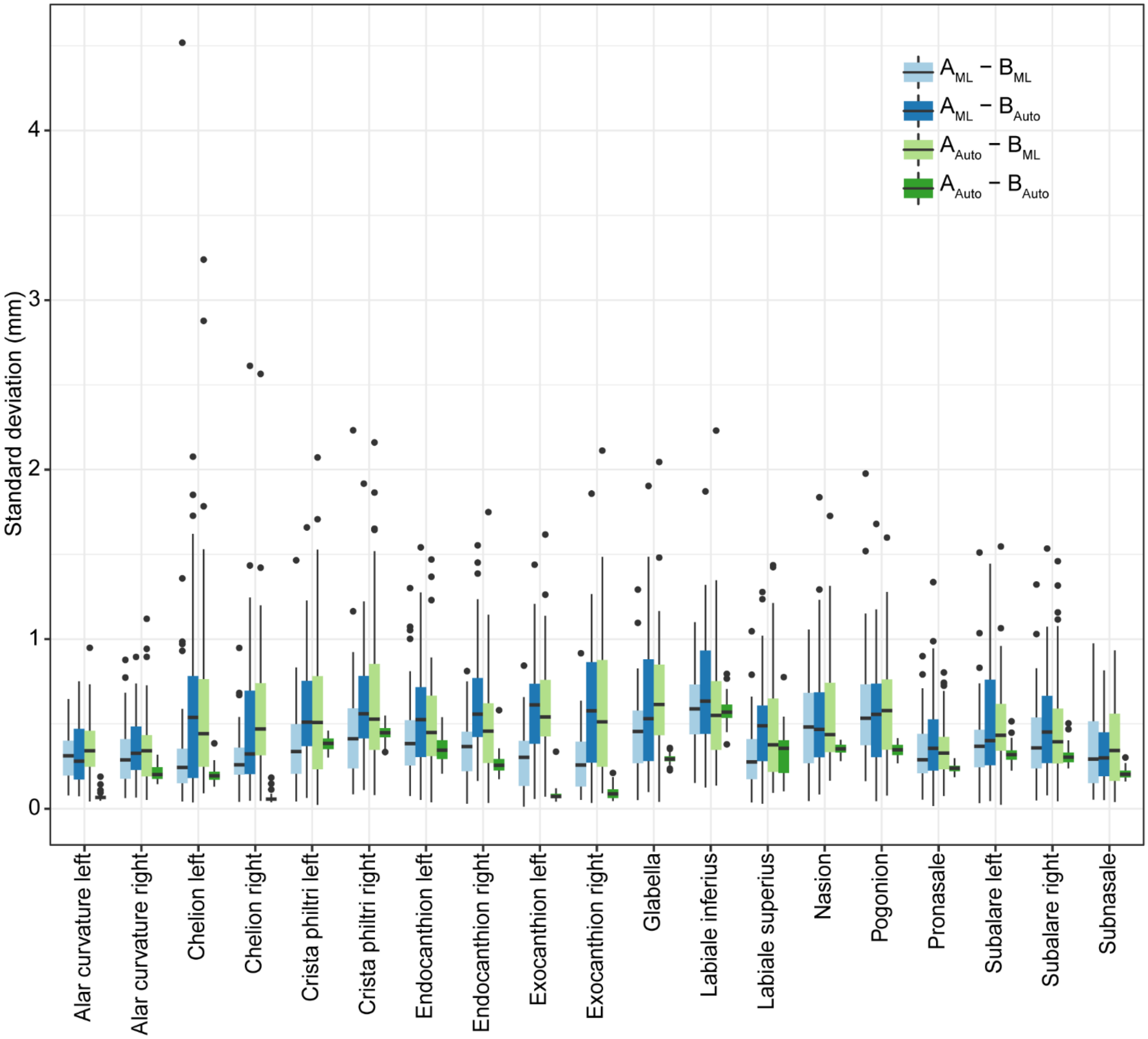


**Supplementary Figure S3. Comparison of inter-observer errors.** Standard deviation values calculated using both manual landmarks and after replacing each observer's set iteratively with their automatic landmarks. All but the labiale superius landmark had significantly smaller variances in the automatic landmark indication comparison ($A_{Auto}$ vs. $B_{Auto}$).

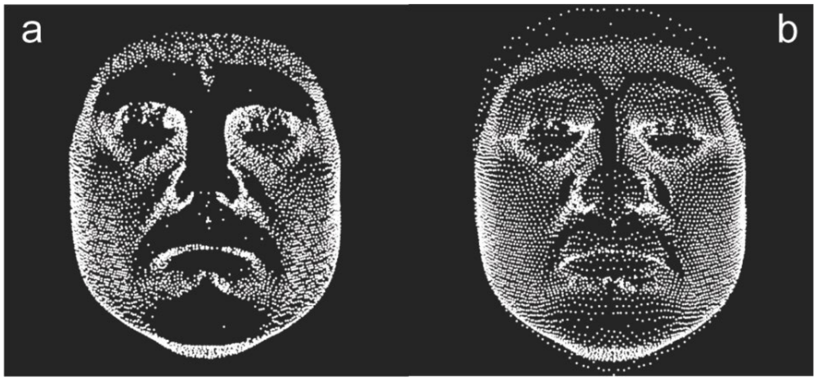


**Supplementary Figure S4. Nonsymmetrical vs. Symmetrical correspondences.** (**a**) Illustration of registration using nonsymmetrical correspondences. (**b**) Illustration of registration using symmetrical correspondences. It can be seen that protruding parts in the target face (e.g. the nose, the front and the chin) are able to co-attract the template in the symmetric setup, therefore the difference in the dark areas in (a) and (b), where no correspondence was found on the template.

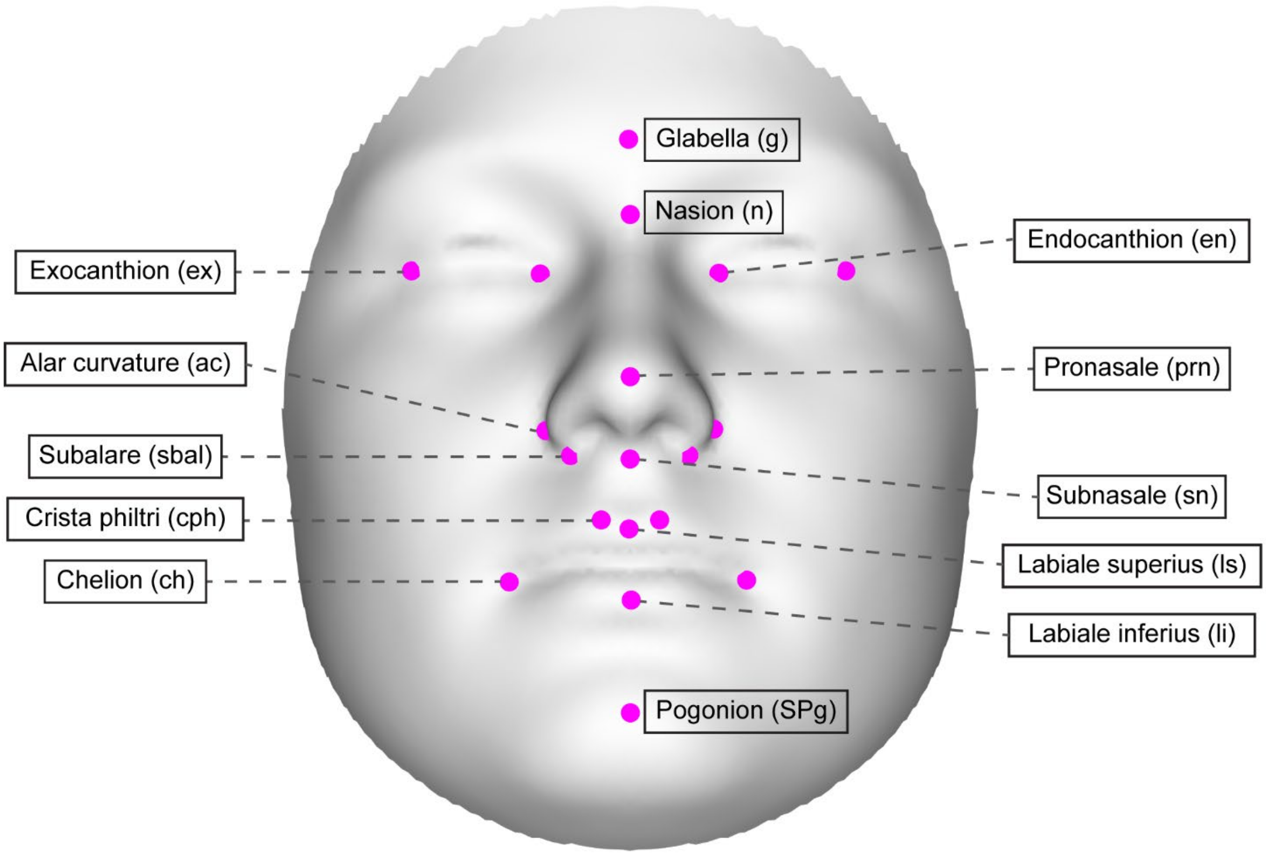


**Supplementary Figure S5. Manual validation landmarks.** Seven midline and twelve bilateral landmarks indicated by two observers during validation of the MeshMonk software. Descriptions of the landmarks are present in Supplementary Table S8.

### *Supplementary Tables*

**Supplementary Table S1.** MANOVAs on average manual landmark configurations and automatic landmark configurations, separately. Results of two separate MANOVAs, one using the average manual landmark configurations from each observer as the response, and the other using the automatic landmark configurations as the response. In both cases, individual and observer were included as predictors. The interaction effect between individual and observer was not included because the residual degrees of freedom became zero when it was included.

| *Variable* | | *DF* | *SS* | *MS* | *R2* | *F* | *Z* | *Pr(>F)* |
|---|---|---|---|---|---|---|---|---|
| *Individual* | ML | 40 | 0.3937 | 0.0098 | 0.9413 | 23.987 | 22.515 | 0.001 |
| | Auto | 40 | 0.3152 | 0.0079 | 0.9714 | 435.70 | 27.609 | 0.001 |
| *Observer* | ML | 1 | 0.0082 | 0.0081 | 0.0195 | 19.853 | 11.563 | 0.001 |
| | Auto | 1 | 0.0085 | 0.0085 | 0.0264 | 472.98 | 8.1969 | 0.001 |
| *Residuals* | ML | 40 | 0.0164 | 0.0004 | 0.0392 | | | |
| | Auto | 40 | 0.0007 | $1.81 \times 10^{-5}$ | 0.0022 | | | |
| *Total* | ML | 81 | 0.4182 | | | | | |
| | Auto | 81 | 0.3245 | | | | | |

**Supplementary Table S2.** Intra- and inter-observer error for the manual landmark indications along the x, y, and z axis, averaged across images for each landmark. Values are standard deviations measured in mm.

| *Landmarks* | *Observer A* | | | | *Observer B* | | | | *Inter-observer* | | | |
|---|---|---|---|---|---|---|---|---|---|---|---|---|
| | *X* | *Y* | *Z* | *Mean* | *X* | *Y* | *Z* | *Mean* | *X* | *Y* | *Z* | *Mean* |
| *Alar curvature left* | 0.21 | 0.68 | 0.92 | 0.60 | 0.15 | 0.55 | 0.61 | 0.43 | 0.12 | 0.38 | 0.42 | 0.31 |
| *Alar curvature right* | 0.21 | 0.75 | 0.93 | 0.63 | 0.13 | 0.46 | 0.54 | 0.38 | 0.12 | 0.34 | 0.53 | 0.33 |
| *Chelion left* | 0.86 | 0.54 | 0.43 | 0.61 | 0.67 | 0.35 | 0.32 | 0.45 | 0.59 | 0.30 | 0.36 | 0.42 |
| *Chelion right* | 0.81 | 0.49 | 0.50 | 0.60 | 0.74 | 0.35 | 0.39 | 0.49 | 0.43 | 0.27 | 0.20 | 0.30 |
| *Crista philtri left* | 0.55 | 0.67 | 0.29 | 0.50 | 0.43 | 0.30 | 0.18 | 0.30 | 0.52 | 0.46 | 0.19 | 0.39 |
| *Crista philtri right* | 0.55 | 0.71 | 0.34 | 0.54 | 0.47 | 0.25 | 0.17 | 0.29 | 0.65 | 0.57 | 0.20 | 0.47 |
| *Endocanthion left* | 0.92 | 0.63 | 0.69 | 0.74 | 0.55 | 0.39 | 0.37 | 0.44 | 0.59 | 0.33 | 0.39 | 0.44 |
| *Endocanthion right* | 1.19 | 0.50 | 0.62 | 0.77 | 0.56 | 0.36 | 0.41 | 0.45 | 0.48 | 0.27 | 0.33 | 0.36 |
| *Exocanthion left* | 0.69 | 0.51 | 0.56 | 0.59 | 0.50 | 0.40 | 0.41 | 0.44 | 0.36 | 0.27 | 0.25 | 0.29 |
| *Exocanthion right* | 0.74 | 0.59 | 0.63 | 0.65 | 0.43 | 0.29 | 0.35 | 0.36 | 0.35 | 0.24 | 0.27 | 0.29 |
| *Glabella* | 0.56 | 0.87 | 0.30 | 0.58 | 0.45 | 1.17 | 0.44 | 0.69 | 0.42 | 0.69 | 0.25 | 0.45 |
| *Labiale inferius* | 0.52 | 0.61 | 0.38 | 0.50 | 0.42 | 0.34 | 0.20 | 0.32 | 0.54 | 0.85 | 0.37 | 0.59 |
| *Labiale superius* | 0.47 | 0.59 | 0.22 | 0.43 | 0.30 | 0.38 | 0.13 | 0.27 | 0.38 | 0.45 | 0.13 | 0.32 |
| *Nasion* | 0.33 | 0.93 | 0.35 | 0.54 | 0.31 | 0.85 | 0.46 | 0.54 | 0.39 | 0.82 | 0.28 | 0.49 |
| *Pogonion* | 0.65 | 1.27 | 0.54 | 0.82 | 0.62 | 1.16 | 0.50 | 0.76 | 0.71 | 0.76 | 0.33 | 0.60 |
| *Pronasale* | 0.42 | 0.71 | 0.25 | 0.46 | 0.30 | 0.44 | 0.21 | 0.32 | 0.29 | 0.49 | 0.21 | 0.33 |
| *Subalare left* | 0.55 | 0.36 | 0.59 | 0.50 | 0.52 | 0.31 | 0.46 | 0.43 | 0.48 | 0.26 | 0.47 | 0.40 |
| *Subalare right* | 0.58 | 0.31 | 0.57 | 0.49 | 0.64 | 0.32 | 0.49 | 0.48 | 0.50 | 0.31 | 0.47 | 0.43 |
| *Subnasale* | 0.38 | 0.65 | 0.32 | 0.45 | 0.33 | 0.72 | 0.35 | 0.47 | 0.18 | 0.55 | 0.31 | 0.35 |
| *Mean* | 0.59 | 0.65 | 0.50 | 0.58 | 0.45 | 0.49 | 0.37 | 0.44 | 0.43 | 0.45 | 0.31 | 0.40 |

**Supplementary Table S3.** MANOVA on all manual landmark indications to assess manual landmarking error. MANOVA was performed on the six GPA-aligned manual landmark indications, with individual, observer, individual x observer, and the nested interaction of observer x iteration as predictors.

| *Variable* | *DF* | *SS* | *MS* | *R2* | *F* | *Z* | *Pr(>F)* |
|---|---|---|---|---|---|---|---|
| *Individual* | 40 | 1.1803 | 0.0295 | 0.8491 | 40.7135 | 26.620 | 0.001 |
| *Observer* | 1 | 0.0244 | 0.0244 | 0.0176 | 33.6563 | 14.568 | 0.001 |
| *Individual x Observer* | 40 | 0.0492 | 0.0012 | 0.0354 | 1.6963 | 26.292 | 0.001 |
| *Observer x Iteration* | 4 | 0.0203 | 0.0051 | 0.0146 | 6.9974 | 19.485 | 0.001 |
| *Residuals* | 160 | 0.1160 | 0.0007 | 0.0834 | | | |
| *Total* | 245 | 1.3901 | | | | | |

**Supplementary Table S4.** Inter-observer error along the x, y, and z axis, averaged across images for each landmark, calculated as the standard deviation (mm) of both manual landmark observers ($A_{ML}$ vs. $B_{ML}$), and after replacing each observer's manual indications with automatic indications ($A_{Auto}$ vs. $B_{ML}$ and $A_{ML}$ vs. $B_{Auto}$). The means for each comparison are also reported in Supplementary Table S5.

| *Landmark* | *$A_{ML}$ vs. $B_{ML}$* | | | | *$A_{ML}$ vs. $B_{Auto}$* | | | | *$A_{Auto}$ vs. $B_{ML}$* | | | |
|---|---|---|---|---|---|---|---|---|---|---|---|---|
| | X | Y | Z | Mean | X | Y | Z | Mean | X | Y | Z | Mean |
| *Alar curvature left* | 0.12 | 0.38 | 0.42 | 0.31 | 0.12 | 0.39 | 0.42 | 0.31 | 0.13 | 0.45 | 0.52 | 0.37 |
| *Alar curvature right* | 0.12 | 0.34 | 0.53 | 0.33 | 0.13 | 0.34 | 0.64 | 0.37 | 0.15 | 0.46 | 0.51 | 0.37 |
| *Chelion left* | 0.59 | 0.30 | 0.36 | 0.42 | 0.84 | 0.54 | 0.50 | 0.63 | 0.97 | 0.49 | 0.46 | 0.64 |
| *Chelion right* | 0.43 | 0.27 | 0.20 | 0.30 | 0.65 | 0.51 | 0.38 | 0.51 | 0.82 | 0.44 | 0.43 | 0.56 |
| *Crista philtri left* | 0.52 | 0.46 | 0.19 | 0.39 | 0.65 | 0.79 | 0.35 | 0.60 | 0.68 | 0.73 | 0.36 | 0.59 |
| *Crista philtri right* | 0.65 | 0.57 | 0.20 | 0.47 | 0.65 | 0.86 | 0.36 | 0.62 | 0.80 | 0.85 | 0.37 | 0.67 |
| *Endocanthion left* | 0.59 | 0.33 | 0.39 | 0.44 | 0.79 | 0.48 | 0.39 | 0.55 | 0.69 | 0.42 | 0.46 | 0.52 |
| *Endocanthion right* | 0.48 | 0.27 | 0.33 | 0.36 | 0.84 | 0.57 | 0.51 | 0.64 | 0.80 | 0.39 | 0.35 | 0.51 |
| *Exocanthion left* | 0.36 | 0.27 | 0.25 | 0.29 | 0.65 | 0.58 | 0.63 | 0.62 | 0.68 | 0.51 | 0.65 | 0.61 |
| *Exocanthion right* | 0.35 | 0.24 | 0.27 | 0.29 | 0.67 | 0.47 | 0.66 | 0.60 | 0.69 | 0.49 | 0.69 | 0.62 |
| *Glabella* | 0.42 | 0.69 | 0.25 | 0.45 | 0.45 | 1.00 | 0.46 | 0.64 | 0.47 | 1.10 | 0.40 | 0.66 |
| *Labiale inferius* | 0.54 | 0.85 | 0.37 | 0.59 | 0.54 | 1.03 | 0.51 | 0.69 | 0.56 | 0.89 | 0.40 | 0.62 |
| *Labiale superius* | 0.38 | 0.45 | 0.13 | 0.32 | 0.49 | 0.70 | 0.30 | 0.50 | 0.52 | 0.72 | 0.24 | 0.49 |
| *Nasion* | 0.39 | 0.82 | 0.28 | 0.49 | 0.38 | 0.89 | 0.38 | 0.55 | 0.39 | 0.93 | 0.39 | 0.57 |
| *Pogonion* | 0.71 | 0.76 | 0.33 | 0.60 | 0.65 | 0.78 | 0.34 | 0.59 | 0.68 | 0.85 | 0.30 | 0.61 |
| *Pronasale* | 0.29 | 0.49 | 0.21 | 0.33 | 0.37 | 0.57 | 0.30 | 0.41 | 0.36 | 0.52 | 0.19 | 0.36 |
| *Subalare left* | 0.48 | 0.26 | 0.47 | 0.40 | 0.62 | 0.38 | 0.44 | 0.48 | 0.63 | 0.33 | 0.58 | 0.51 |
| *Subalare right* | 0.50 | 0.31 | 0.47 | 0.43 | 0.65 | 0.38 | 0.53 | 0.52 | 0.53 | 0.37 | 0.53 | 0.47 |
| *Subnasale* | 0.18 | 0.55 | 0.31 | 0.35 | 0.25 | 0.47 | 0.24 | 0.32 | 0.25 | 0.54 | 0.29 | 0.36 |
| *Mean* | 0.43 | 0.45 | 0.31 | 0.40 | 0.55 | 0.62 | 0.44 | 0.53 | 0.57 | 0.60 | 0.43 | 0.53 |

**Supplementary Table S5.** Comparison of inter-observer errors. Standard deviation for only manual landmarks and for manual and automatic comparisons. Based on a paired T-test, comparisons that are significantly different using an alpha of 0.05 are in bold.

| | $A_{ML}$-$B_{ML}$ | $A_{ML}$ – $B_{Auto}$ | | | $A_{Auto}$ - $B_{ML}$ | | |
|---|---|---|---|---|---|---|---|
| *Landmark* | *Mean SD (mm)* | *Mean SD (mm)* | *T statistic* | *P value* | *Mean SD (mm)* | *T statistic* | *P value* |
| *Alar curvature left* | 0.31 | 0.31 | -0.10 | 0.9197 | 0.37 | -2.14 | **0.0382** |
| *Alar curvature right* | 0.33 | 0.37 | -0.99 | 0.3275 | 0.37 | -1.23 | 0.2266 |
| *Chelion left* | 0.42 | 0.63 | -1.89 | 0.0665 | 0.64 | -2.40 | **0.0212** |
| *Chelion right* | 0.30 | 0.51 | -2.59 | **0.0132** | 0.56 | -3.78 | **$5.20 \times 10^{-4}$** |
| *Crista philtri left* | 0.39 | 0.60 | -2.87 | **0.0066** | 0.59 | -3.11 | **0.0034** |
| *Crista philtri right* | 0.47 | 0.62 | -1.89 | 0.0661 | 0.67 | -3.53 | **0.0010** |
| *Endocanthion left* | 0.44 | 0.55 | -2.50 | **0.0167** | 0.52 | -1.46 | 0.1527 |
| *Endocanthion right* | 0.36 | 0.64 | -5.49 | **$2.50 \times 10^{-6}$** | 0.51 | -2.84 | **0.0071** |
| *Exocanthion left* | 0.29 | 0.62 | -6.14 | **$3.00 \times 10^{-7}$** | 0.61 | -5.80 | **$8.93 \times 10^{-7}$** |
| *Exocanthion right* | 0.29 | 0.60 | -5.54 | **$2.09 \times 10^{-6}$** | 0.62 | -5.65 | **$1.49 \times 10^{-6}$** |
| *Glabella* | 0.45 | 0.64 | -2.63 | **0.0121** | 0.66 | -3.12 | **0.0033** |
| *Labiale inferius* | 0.59 | 0.69 | -2.14 | **0.0381** | 0.62 | -0.54 | 0.5895 |
| *Labiale superius* | 0.32 | 0.50 | -3.05 | **0.0040** | 0.49 | -3.21 | **0.0026** |
| *Nasion* | 0.49 | 0.55 | -0.93 | 0.3556 | 0.57 | -1.25 | 0.2188 |
| *Pogonion* | 0.60 | 0.59 | 0.15 | 0.8835 | 0.61 | -0.22 | 0.8245 |
| *Pronasale* | 0.33 | 0.41 | -2.16 | **0.0366** | 0.36 | -0.65 | 0.5169 |
| *Subalare left* | 0.40 | 0.48 | -1.35 | 0.1842 | 0.51 | -2.24 | **0.0308** |
| *Subalare right* | 0.43 | 0.52 | -1.63 | 0.1114 | 0.47 | -0.84 | 0.4083 |
| *Subnasale* | 0.35 | 0.32 | 0.69 | 0.4939 | 0.36 | -0.26 | 0.7930 |
| *Mean* | 0.40 | 0.53 | | | 0.53 | | |

**Supplementary Table S6.** Comparison of error variance. The standard deviation of average landmark configurations for the manual ($A_{ML}$ vs. $B_{ML}$) and automatic ($A_{Auto}$ vs. $B_{Auto}$) landmarks, averaged across scans. Levene's test was performed per landmark to assess the difference between error variance and comparisons that are significantly different using an alpha of 0.05 are in bold.

| *Landmark* | *Manual (mm)* | *Auto (mm)* | *F value* | *P value* |
|---|---|---|---|---|
| *Alar curvature left* | 0.3067 | 0.0728 | 59.6244 | **2.83 x $10^{-11}$** |
| *Alar curvature right* | 0.3287 | 0.2133 | 22.2346 | **1.01 x $10^{-5}$** |
| *Chelion left* | 0.4182 | 0.1998 | 4.6453 | **0.0341** |
| *Chelion right* | 0.2984 | 0.0637 | 24.5101 | **4.03 x $10^{-6}$** |
| *Crista philtri left* | 0.3881 | 0.3811 | 29.1832 | **6.60 x $10^{-7}$** |
| *Crista philtri right* | 0.4737 | 0.4472 | 18.1685 | **5.49 x $10^{-5}$** |
| *Endocanthion left* | 0.4362 | 0.3504 | 14.2000 | **3.13 x $10^{-4}$** |
| *Endocanthion right* | 0.3608 | 0.2669 | 28.4103 | **8.85 x $10^{-7}$** |
| *Exocanthion left* | 0.2946 | 0.0808 | 47.7334 | **1.06 x $10^{-9}$** |
| *Exocanthion right* | 0.2855 | 0.0961 | 28.0100 | **1.03 x $10^{-6}$** |
| *Glabella* | 0.4542 | 0.2938 | 41.5866 | **7.95 x $10^{-9}$** |
| *Labiale inferius* | 0.5857 | 0.5773 | 26.3847 | **1.93 x $10^{-6}$** |
| *Labiale superius* | 0.3185 | 0.3289 | 2.4213 | 0.1236 |
| *Nasion* | 0.4938 | 0.3511 | 87.7550 | **1.67 x $10^{-14}$** |
| *Pogonion* | 0.5987 | 0.3478 | 23.9927 | **4.95 x $10^{-6}$** |
| *Pronasale* | 0.3323 | 0.2376 | 38.2428 | **2.49 x $10^{-8}$** |
| *Subalare left* | 0.4005 | 0.3239 | 16.4805 | **1.14 x $10^{-4}$** |
| *Subalare right* | 0.4283 | 0.3113 | 25.6819 | **2.54 x $10^{-6}$** |
| *Subnasale* | 0.3480 | 0.2072 | 42.6476 | **5.57 x $10^{-9}$** |
| *Mean* | 0.3974 | 0.2711 | | |

**Supplementary Table S7.** Descriptive data for the validation sample used. These data are included only to describe the variation present in the sample and were not used as covariates in statistical analyses.

| *ID* | *Sex* | *Age* | *Height (cm)* | *Weight (kg)* | *Self-identified population* | *Year, Camera* |
|---|---|---|---|---|---|---|
| *MM_124* | F | 33 | 155.00 | 48.00 | Admixed African and European | 2006, 2-pod |
| *MM_172* | F | 20 | 158.00 | 43.00 | Italian | 2006, 2-pod |
| *MM_130* | M | 21 | 175.00 | 71.99 | Italian | 2006, 2-pod |
| *MM_154* | M | 22 | 178.00 | 60.01 | Polish | 2006, 2-pod |
| *MM_156* | M | 21 | 172.00 | 57.02 | Portuguese | 2006, 2-pod |
| *MM_143* | M | 41 | 176.00 | 76.02 | Portuguese | 2006, 2-pod |
| *MM_137* | F | 28 | 169.00 | 57.02 | Portuguese | 2006, 2-pod |
| *MM_150* | M | 24 | 175.00 | 65.00 | Portuguese | 2006, 2-pod |
| *MM_153* | F | 30 | 165.00 | 66.00 | Portuguese | 2006, 2-pod |
| *MM_179* | F | 20 | 165.10 | 57.15 | Irish | 2006, 2-pod |
| *MM_188* | F | 27 | 165.00 | 57.15 | Irish | 2006, 2-pod |
| *MM_142* | F | 43 | 165.10 | 69.85 | Irish | 2006, 2-pod |
| *MM_155* | M | 19 | 184.00 | 60.33 | Irish | 2006, 2-pod |
| *MM_125* | M | 19 | 180.00 | 66.00 | Irish | 2006, 2-pod |
| *MM_125* | M | 23 | 172.55 | 72.85 | Broadly European | 2013, 2-pod |
| *MM_175* | F | 33 | 160.70 | 57.40 | Unknown | 2014, 3-pod |
| *MM_143* | M | 19 | 179.00 | 103.80 | Mixed European and European American | 2014, 3-pod |
| *MM_165* | M | 62 | 166.00 | 63.30 | Mixed European and European American | 2014, 3-pod |
| *MM_147* | F | 26 | 165.00 | 101.30 | Mixed European and European American | 2014, 3-pod |
| *MM_102* | F | 73 | 169.00 | 83.00 | Mixed European and European American | 2014, 3-pod |
| *MM_189* | M | 21 | 168.60 | 70.80 | Broadly European | 2014, 3-pod |
| *MM_147* | F | 79 | 149.86 | 70.31 | Broadly European | 2014, 3-pod |
| *MM_148* | F | 66 | 170.18 | 97.07 | Mixed European and European American | 2014, 3-pod |
| *MM_142* | F | 24 | 154.94 | 52.16 | Mixed European and European American | 2014, 3-pod |
| *MM_175* | F | 63 | 170.18 | 81.65 | Broadly European | 2014, 3-pod |
| *MM_170* | F | 53 | 162.56 | 86.18 | Broadly European | 2014, 3-pod |
| *MM_157* | F | 25 | 152.40 | 46.72 | Mixed European and European American | 2014, 3-pod |
| *MM_157* | F | 55 | 165.10 | 62.60 | Mixed European and European American | 2014, 3-pod |
| *MM_177* | F | 20 | 162.56 | 68.04 | Mixed European and European American | 2014, 3-pod |
| *MM_138* | F | 65 | 157.48 | 54.43 | Mixed European and European American | 2014, 3-pod |
| *MM_182* | F | 27 | 156.40 | 62.40 | Mixed European and European American | 2014, 3-pod |
| *MM_120* | F | 19 | 181.00 | 68.40 | Mixed European and European American | 2014, 3-pod |
| *MM_173* | F | 18 | 179.00 | 65.00 | Broadly European | 2014, 3-pod |
| *MM_186* | F | 20 | 162.30 | 98.90 | Broadly European | 2014, 3-pod |
| *MM_132* | F | 19 | 170.00 | 75.70 | Broadly European | 2014, 3-pod |
| *MM_174* | F | 53 | 164.00 | 50.80 | Mixed European, Jewish | 2014, 3-pod |
| *MM_200* | F | 22 | 169.90 | 55.00 | Mixed European and European American | 2014, 3-pod |
| *MM_160* | M | 28 | 174.00 | 65.30 | Broadly European | 2014, 3-pod |
| *MM_193* | F | 23 | 161.00 | 78.20 | Broadly European | 2014, 3-pod |
| *MM_171* | F | 19 | 162.00 | 59.40 | Mixed European, Jewish | 2014, 3-pod |
| *MM_141* | F | 21 | 164.60 | 67.20 | Mixed European and European American | 2014, 3-pod |

**Supplementary Table S8.** Description of landmarks used in validation. Landmark descriptions are those reported on the Richtsmeier Lab website (http://www.getahead.la.psu.edu/).

| *Landmark* | *Abbr.* | *Location* | *Definition* |
|---|---|---|---|
| *Glabella* | g | Midline | The most prominent midline point between the eyebrows. |
| *Nasion* | n | Midline | The point in the midline of both the nasal root and the nasofrontal suture. This point is always above the line that connects the two inner canthi. |
| *Pronasale* | prn | Midline | The most protruded point of the apex nasi. |
| *Subnasale* | sn | Midline | The midpoint of the angle at the columella base where the lower border of the nasal septum and the surface of the upper lip meet. |
| *Labiale superius* | ls | Midline | The midpoint of the upper vermillion line. |
| *Labiale inferius* | li | Midline | The midpoint of the lower vermillion line. |
| *Pogonion* | SPg | Midline | The most anterior point of the chin. |
| *Endocanthion* | en | Bilateral | The point at the inner commissure of the eye fissure. |
| *Exocanthion* | ex | Bilateral | The point at the outer commissure of the eye fissure. |
| *Alar curvature* | ac | Bilateral | The most lateral point in the curved base of each ala. Indicating the facial insertion of the nasal wingbase. |
| *Subalare* | sbal | Bilateral | The point at the lower limit of each alar base, where the alar base disappears into the skin of the upper lip. The landmarks indicate the labial insertion of the alar base |
| *Crista philtri* | cph | Bilateral | The lower point on each elevated margin of the philtrum just above the vermillion line. |
| *Chelion* | ch | Bilateral | Point located at each labial commissure at the most lateral intersection of upper and lower lip. |

### ***Supplementary Methods***

#### **MeshMonk Registration process**

<u>Finding correspondences</u>

The symmetrical correspondences are calculated by combining two affinity matrices: 1) from template points to points on the target surface (push forces; the typical one-to-one correspondences calculation), and 2) from target points to points on the template surface (pull forces). This ensures that potential protrusions present on the target surface are allowed to pull the corresponding structure on the template, as illustrated in Supplementary Fig. S4C. Binary correspondences are avoided by using the weighted k-nearest neighbor rule, allowing correspondence to be defined as an interpolation between existing surface points (anywhere on the surface). For each point, the k closest points on the opposite surface are searched for, and the inverse of the distance to each of its closest points is coded as a weight in the affinity matrix. The weights for the remaining non-closest points are set to zero. The distance to the closest points can be computed in terms of 3D position only or a combination of 3D position and 3D normal orientation in each point, rendering 6D distances as a definition for “closeness”. The incorporation of the normal information better matches points with a similar orientation and avoids the inappropriate matching of opposite oriented points. For example, in skeletal surface data, the inner and outer surface have opposing normal orientations and the left and right flanks of the human nose also have opposing normal orientations.

<u>Pruning correspondences</u>

The MeshMonk toolbox allows for the identification of outliers either deterministically or stochastically, or a combination of both. Deterministic outliers include correspondences to surface border points (this properly handles the situation when the template and target surface have non-overlapping structures), large triangles (as identified using a z-score on triangle area, and therefore defined in function of the underlying surface resolution), angle between point surface normals (further excluding point correspondences with opposing normal orientations) and any manually tagged points on the target or template. Stochastic outliers are defined following[1], using inlier versus outlier distribution estimations. The inlier distances are assumed to form a Gaussian distribution, and any point falling out of +/- a user-defined κ times the standard deviation is considered abnormal and flagged as an outlier. Then, the contributions of the outlier correspondences are fixed and the confidence values of all the points are updated.

**MeshMonk Parameter Tuning**

The accuracy of the template registration can be determined by the "shape fit" defined by the root mean squared distance (RMSD) of all template points to the target surface after registration, and the quality of the shape model based on the minimum description length. For this, a principal component analysis (PCA) can be performed, where the amount of variance of the model is explained in terms of the number of principal components (PCs). Intuitively, the smaller the RMSD and the lesser PCs (or the more variance explained with a fixed number of PCs) required to explain the model, the better the registration. Hereto, in order to improve the robustness of MeshMonk for the facial image registration used in this work, three main parameters were tuned: a) the number of iterations for the non-rigid registration, b) the number of neighbors involved in the transformation's regularization, and c) the number of k-nearest neighbors used to find the best correspondences. For each, a range of values was tested and the respective RMSD and PCA were calculated.

<u>Number of iterations for the non-rigid registration</u>
Typically, the fidelity of the template registration comes at the expenses of computational time. The higher the number of iterations for the non-rigid registration, the better the template registration, but also the higher the computational time. This is especially of concern when registering a large dataset, which might jeopardize the efficiency of the process. Therefore, a tradeoff between the number of iterations and the computational time has to be made. Here, the accuracy of the registration was calculated using 10 to 300 iterations. As shown in Supplementary Fig. S6A, the higher the number of iterations the smaller the RMSD (i.e. the better the shape fit). However, it can also be seen that after about 150 iterations the amount of variance explained in the top 10 PCs becomes stable. Therefore, in seeking computational efficiency, a good compromise between computational time and accuracy can be obtained by selecting about 200 iterations.

<u>Number of neighbors regularization</u>
In theory, the amount of smoothness of the transformation is controlled by iteratively convolving the estimated displacement field with a narrower Gaussian each time, in this way narrowing the scope from global to local. However, in practice, when the number of template nodes is very large this procedure becomes computationally expensive. Thus, an implementation can be done where instead of convolving the global transformation with a narrower Gaussian in every iteration, a fixed width is chosen and a small number of neighbors (n) is smoothed iteratively. As shown in Supplementary Fig. S6B, the accuracy of the registration was calculated based on n=10 to n=160 neighbors. In this case, it can be seen that for both quality measurements, a plateau is reach at about n=70 neighbors. Since no improvement is expected by using more neighbors, a conservative measure is achieved by selecting about 80 neighbors. It is also observed than an increased amount of smoothing by enlarging the number of neighbors increases the error on "shape fit." This is not expected, as any kind of smoothing encourages points to exactly map onto the target surface.

However, the increase in error is marginal, and the final RMSE value at 80 neighbors is still below one tenth of a millimeter.

Number of k-nearest neighbors

MeshMonk uses the weighted k-nearest neighbor rule to assign correspondences based on the interpolation between existing surface points. The precision of the correspondences is directly influenced by the position of the chosen nearest points. To avoid misleading assignments, the question: “how near is near?” needs to be answered. Therefore, the accuracy of the template registration was calculated using from 3 to 20 nearest neighbors and the results are shown in Supplementary Fig. S6C. Contrary to the previous parameters, in this case the smaller the number of k-nearest neighbors the better the registration. As can be intuitively induced, the best combination to find the corresponding average are the three nearest points, any point outside of it might deviate the response, as can be seen from both the shape and the model registration.

In conclusion, in order to ensure a proper surface registration while limiting computational time 200 non-rigid iterations, 80 neighbors for the regularization of the transformation and k=3 nearest points for finding the correspondences on a dataset of 3D facial images are the best combination.

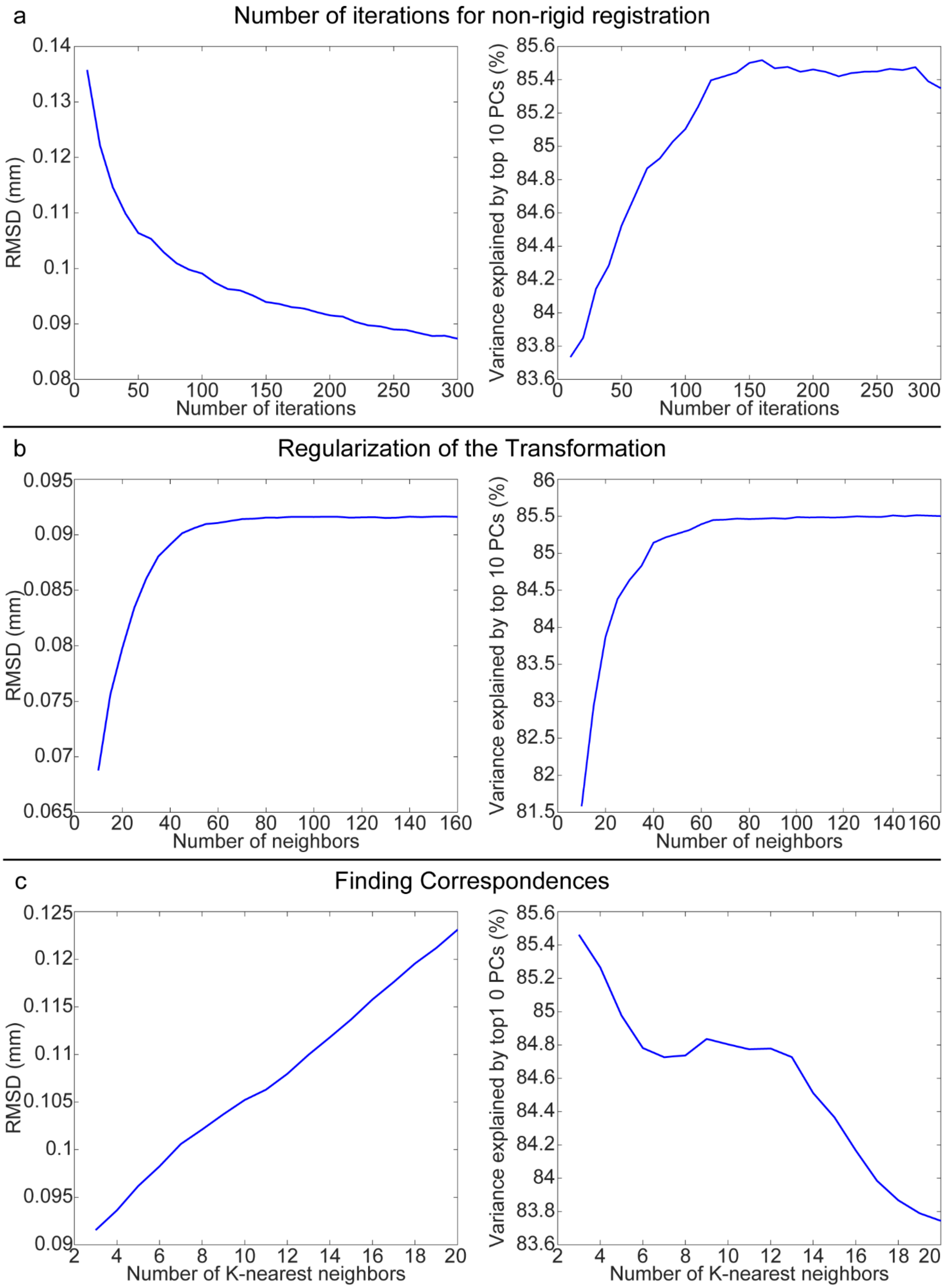


**Supplementary Figure S6.** Parameter tuning of the template registration calculated based on (**a**) the number of iterations for the non-rigid registration, (**b**) the number of neighbors for the transformation regularization and (**c**) the number of k-nearest neighbor to find the correspondences.

### Automatic placement of validation landmarks

Performing the automatic landmark indications requires the transfer of the manual landmarks of each surface image to the mapped representation of that image. Once in the registration's space, the landmarks of 40 subjects x 2 observers x 3 indications = 240 can be placed onto the anthropometric mask and their average placement can be found. To find the corresponding location of the manual landmarking onto the mapped image, a search for the three closest points is performed and the landmark location is computed as their average. However, this does not ensure that the corresponding point lays on the surface, requiring a barycentric coordinate transformation. Thus, two coordinate transformations are performed, one from Cartesian to Barycentric to establish the corresponding point from the surface image to the registered (mapped using MeshMonk) image, and one from Barycentric to Cartesian to actually place it onto the template surface. Thus, once all the landmark indications are expressed in the registration space, they can be transferred on the original template and their average can be found. Now, to automatically superimpose the average landmarks indications on the left-out image, the reversed coordinate transformation is performed, this time going from the template image to the left-out image. A brief pseudo-code description of the process is as follows:

**Input:** 41 images x 6 landmark indications
**Output:** Automatic landmarking placement
**Procedure:**
1. ***for*** *i* = 1 to 41:
1.1 Image registration
1.2. Manual landmarks aligned by Cartesian to Barycentric coordinate transformation from the manually landmarked image to the mapped surface
    ***end***
2. ***for*** *j* = 1 to 40:
Manual landmarks placed directly on the mapped surface by Barycentric to Cartesian coordinate transformation to transfer all the landmarks to the template
    ***end***
3. Average the landmarking positions on the template
4. For the left-out image:
4.1. Automatic Landmarks aligned by Cartesian to Barycentric coordinate transformation from the template to the unmapped left-out image
4.2. Automatic Landmarks placed on the unmapped image by Barycentric to Cartesian coordinate transformation
5. Repeat steps 2 through 4 until all N=41 images have been automatically landmarked.
6. ***Done***
